## Supplementary material for "Pleiotropic roles for mycobacterial DinB2 in frameshift and substitution mutagenesis": SI Tables and Figs

### Supplementary tables

| Strains | Genetic background | reference or source |
|---|---|---|
| *M. smegmatis* | | |
| mc$^2$155 | Wild type | (1) |
| PDS353 | Δ*recA* | (2) |
| PDS139 | Δ*dnaE2* | (2) |
| Mgm4062 | pmsg419 | This work |
| mgm4063 | pRGM47 | This work |
| mgm4072 | pRGM48 | This work |
| mgm4073 | pRGM49 | This work |
| mgm4074 | pRGM50 | This work |
| PDS416 | pDP69 | This work |
| PDS417 | pDP70 | This work |
| *E.coli* | | |
| DH5α | F$^-$ Φ80*lac*ZΔM15 Δ(*lac*ZYA-*arg*F) U169 *rec*A1 *end*A1 *hsd*R17($r_k^-$, $m_k^+$) *pho*A *sup*E44 *thi*-1 *gyr*A96 *rel*A1 $\lambda^-$ | Lab collection |

Table S1: strains used in this study

| Plasmids | description | Cloning enzyme sites | Cloning primers (infusion reaction) | References or sources |
|---|---|---|---|---|
| pmsg419 | ATc-on system vector (hyg$^{R}$, OriMyc) | | | Lab Stock |
| pDB60 | Mycob. integr. vector (Strep$^{R}$, attP(L5)) | | | Lab Stock |
| pRGM47 | pmsg419-*dinB2* Strep tag | *ClaI* | dinB2fw-dinB2rev1 | This work |
| pRGM48 | pmsg419-*dinB2* | *ClaI* | dinB2fw-dinB2rev2 | This work |
| pRGM49 | pmsg419-*dinB2*$^{D107A}$ Strep tag | *ClaI* | dinB2fw-dinB2$^{cat}$rev+dinB2$^{cat}$fw-dinB2rev1 | This work |
| pRGM50 | pmsg419-*dinB2*$^{L14F}$ Strep tag | *ClaI* | dinB2fw-dinB2$^{steric}$rev+dinB2$^{steric}$fw-dinB2rev1 | This work |
| pDP69 | pmsg419-*dinB3* | *ClaI* | ODP197-ODP298 | This work |
| pDP70 | pmsg419-*dinB3* Strep tag | *ClaI* | ODP197-ODP299 | This work |
| pDP120 | pDB60 derivative with *kan*::3T | | | (3) |
| pDP121 | pDB60 derivative with *kan*::3C | | | (3) |
| pDP122 | pDB60 derivative with *kan*::3G | | | (3) |
| pDP123 | pDB60 derivative with *kan*::3A | | | (3) |
| pDP124 | pDB60 derivative with *kan*::4T | | | (3) |
| pDP125 | pDB60 derivative with *kan*::4C | | | (3) |
| pDP126 | pDB60 derivative with *kan*::4G | | | (3) |
| pDP127 | pDB60 derivative with *kan*::4A | | | (3) |
| pDP128 | pDB60 derivative with *kan*::6T | | | (3) |
| pDP129 | pDB60 derivative with *kan*::6C | | | (3) |
| pDP130 | pDB60 derivative with *kan*::6G | | | (3) |
| pDP131 | pDB60 derivative with *kan*::6A | | | (3) |
| pDP132 | pDB60 derivative with *kan*::7T | *EcoR1* | ODP443-ODP445+ODP458-ODP444 | This work |
| pDP133 | pDB60 derivative with *kan*::7C | *EcoR1* | ODP443-ODP445+ODP459-ODP444 | This work |
| pDP134 | pDB60 derivative with *kan*::7G | *EcoR1* | ODP443-ODP445+ODP460-ODP444 | This work |
| pDP135 | pDB60 derivative with *kan*::7A | *EcoR1* | ODP443-ODP445+ODP461-ODP444 | This work |
| pDP136 | pDB60 derivative with *kan*::9T | *EcoR1* | ODP443-ODP445+ODP462-ODP444 | This work |
| pDP137 | pDB60 derivative with *kan*::9C | *EcoR1* | ODP443-ODP445+ODP463-ODP444 | This work |
| pDP138 | pDB60 derivative with *kan*::9G | *EcoR1* | ODP443-ODP445+ODP464-ODP444 | This work |
| pDP139 | pDB60 derivative with *kan*::9A | *EcoR1* | ODP443-ODP445+ODP465-ODP444 | This work |
| pDP144 | pDB60 derivative with *kan*::5T | | | (3) |
| pDP145 | pDB60 derivative with *kan*::5C | | | (3) |
| pDP146 | pDB60 derivative with *kan*::5G | | | (3) |
| pDP147 | pDB60 derivative with *kan*::5A | | | (3) |
| pDP148 | pDB60 derivative with *kan*::8T | *EcoR1* | ODP443-ODP445+ODP494-ODP444 | This work |
| pDP149 | pDB60 derivative with *kan*::8C | *EcoR1* | ODP443-ODP445+ODP495-ODP444 | This work |
| pDP150 | pDB60 derivative with *kan*::8G | *EcoR1* | ODP443-ODP445+ODP496-ODP444 | This work |
| pDP151 | pDB60 derivative with *kan*::8A | *EcoR1* | ODP443-ODP445+ODP497-ODP444 | This work |
| pDP186 | pDB60 derivative with *sacB*::9C | *EcoR1* | ODP593-ODP596+ODP597-ODP598 | This work |
| pDP194 | pDB60 derivative with *sacB*::6C | *EcoR1* | ODP593-ODP614+ODP615-ODP598 | This work |

Table S2: plasmids used in this study

| Primers | Sequences (5’→3’)* | Targets | Vectors and cloning sites |
|---|---|---|---|
| *dinB*s inducible expression constructs | | | |
| dinB2fw | **CAGAAAGGAGGCCAT**ATGACCAAATGGGTGCTC | fw *dinB2* | pmsg419 (*ClaI*) |
| dinB2rev1 | **AGGTCGACGGTATCG**ATACTACTTTTCGAACTGCGGGTGGCTCCAG GTGCCTGCAGTGACAG | rev *dinB2* + streptavidin tag | pmsg419 (*ClaI*) |
| dinB2rev2 | **AGGTCGACGGTATCG**ATGTGCTCGAGTTAGGTGCCTGCAGTGAC | rev *dinB2* | pmsg419 (*ClaI*) |
| dinB2$^{cat}$rev | **GCCCAGATACGCCTCGGC**CCAGCCCCACACCTCCAAC | rev internal *dinB2*$^{Msm}$ with pol. dead mut. (D107A) | pmsg419 (*ClaI*) |
| dinB2$^{cat}$fw | **GCC**GAGGCGTATCTGGGC | fw internal *dinB2*$^{Msm}$ with pol. dead mut. (D107A) | pmsg419 (*ClaI*) |
| dinB2$^{steric}$rev | **GCAACTCCACCGAAGCAAA**GAACTGGTCCAGATCGAC | rev internal *dinB2*$^{Msm}$ with steric gate mut. (L14F) | pmsg419 (*ClaI*) |
| dinB2$^{steric}$fw | **TTT**GCTTCGGTGGAGTTGC | fw internal *dinB2*$^{Msm}$ with steric gate mut. (L14F) | pmsg419 (*ClaI*) |
| ODP297 | **CAGAAAGGAGGCCAT**ATGTTCGTGTCCGCTGC | fw *dinB3* | pmsg419 (*ClaI*) |
| ODP298 | **AGGTCGACGGTATCG**CTAGTCCGGCAGCATGG | rev *dinB3* | pmsg419 (*ClaI*) |
| ODP299 | **AGGTCGACGGTATCG**CTACTTTTCGAACTGCGGGTGGCTCCAGTCC GGCAGCATGGG | rev *dinB3* + streptavidin tag | pmsg419 (*ClaI*) |
| *kan* inactivated by homo-oligonucleotide runs | | | |
| ODP443 | **TCCAGCTGCAGAATT**TCCCAAGGACACTGAGTCC | fw *kan* | pDB60 (*EcoR1*) |
| ODP444 | **GATAAGCTTCGAATT**TTGCTGACTCATACCAGGC | rev *kan* | pDB60 (*EcoR1*) |
| ODP445 | **CATAACACCCCTTGT**ATTACTG | internal rev *kan* | pDB60 (*EcoR1*) |
| ODP458 | **ACAAGGGGTGTTATG***TTTTTTT*AGCCATATTCAACGGGAAACG | internal fw *kan* (7T addition) | pDB60 (*EcoR1*) |
| ODP459 | **ACAAGGGGTGTTATG***CCCCCCC*AGCCATATTCAACGGGAAACG | internal fw *kan* 7C addition) | pDB60 (*EcoR1*) |
| ODP460 | **ACAAGGGGTGTTATG***GGGGGG*AAGCCATATTCAACGGGAAACG | internal fw *kan* (7G addition) | pDB60 (*EcoR1*) |
| ODP461 | **ACAAGGGGTGTTATG**G*AAAAAAA*GCCATATTCAACGGGAAACG | internal fw *kan* (7A addition) | pDB60 (*EcoR1*) |
| ODP462 | **ACAAGGGGTGTTATG***TTTTTTTTT*AGCCATATTCAACGGGAAACG | internal fw *kan* (9T addition) | pDB60 (*EcoR1*) |
| ODP463 | **ACAAGGGGTGTTATG***CCCCCCCCC*AGCCATATTCAACGGGAAACG | internal fw *kan* (9C addition) | pDB60 (*EcoR1*) |
| ODP464 | **ACAAGGGGTGTTATG***GGGGGGGG*AAGCCATATTCAACGGGAAACG | internal fw *kan* (9G addition) | pDB60 (*EcoR1*) |
| ODP465 | **ACAAGGGGTGTTATG**G*AAAAAAAAA*GCCATATTCAACGGGAAACG | internal fw *kan* (9A addition) | pDB60 (*EcoR1*) |
| ODP494 | **ACAAGGGGTGTTATG***TTTTTTTT*AGCCATATTCAACGGGAAACG | internal fw *kan* (8T addition) | pDB60 (*EcoR1*) |
| ODP495 | **ACAAGGGGTGTTATG***CCCCCCCC*AGCCATATTCAACGGGAAACG | internal fw *kan* (8C addition) | pDB60 (*EcoR1*) |
| ODP496 | **ACAAGGGGTGTTATG***GGGGGGG*AAGCCATATTCAACGGGAAACG | internal fw *kan* (8G addition) | pDB60 (*EcoR1*) |
| ODP497 | **ACAAGGGGTGTTATG**G*AAAAAAAA*GCCATATTCAACGGGAAACG | internal fw *kan* (8A addition) | pDB60 (*EcoR1*) |
| *sacB* inactivated by homo-oligonucleotide runs | | | |
| ODP593 | **TCCAGCTGCAGAATT**AACCCATCACATATACCTGCCG | fw *sacB* | pDB60 (*EcoR1*) |
| ODP596 | **GTT*GGGGGGGGG*CAT**CGTTCATGTCTCCTTTTTTATG | internal rev *sacB* (9C addition) | pDB60 (*EcoR1*) |
| ODP597 | **ATG*CCCCCCCCC*AAC**ATCAAAAAGTTTGCAAAACAAG | internal fw *sacB* (9C addition) | pDB60 (*EcoR1*) |
| ODP598 | **GATAAGCTTCGAATT**ACTATCAATAAGTTGGAGTCATTACC | rev *sacB* | pDB60 (*EcoR1*) |
| ODP614 | **GTT*GGGGGG*CAT**CGTTCATGTCTCCTTTTTTATG | internal rev *sacB* (6C addition) | pDB60 (*EcoR1*) |
| ODP615 | **ATG*CCCCCC*AAC**ATCAAAAAGTTTGCAAAACAAG | internal fw *sacB* (6C addition) | pDB60 (*EcoR1*) |
| Screening PCR and sequencing | | | |
| ODP236 | CTCCCTATCAGTGATAGATAGGCTCTGG | fw PCR screening and seq pmsg419 cloning | |
| ODP237 | CATGACCAACTTCGATAACGTTCTCGG | rev PCR screening and seq pmsg419 cloning | |
| ODP474 | TGATTCTGTGGATAACCGTATTACCGCCTTTG | fw PCR screening and seq pDB60 cloning | |
| ODP475 | AAGGCCCAGTCTTTCGACTGAGC | rev PCR screening and seq pDB60 cloning | |

| | | |
|---|---|---|
| ODP378 | CAAGAAGCTGGGCCTGAACGC | fw *rpoB* PCR |
| ODP379 | GCGGTTGGCGTCGTCGTG | rev *rpoB* PCR |
| ODP380 | GAGCGTGTCGTGCGTGAG | *rpoB* seq |
| ODP476 | TGGCCTTTTGCTGGCCTTTTGC | fw *kan or sacB* PCR |
| ODP477 | TTCAACAAAGCCGCCGTCCC | rev *kan* PCR |
| ODP479 | ACTGAATCCGGTGAGAATGG | *kan* seq |
| ODP172 | TTAGACGTAATGCCGTCAATCGTC | rev *sacB* PCR |
| ODP474 | TGATTCTGTGGATAACCGTATTACCGCCTTTG | *sacB* seq |
| In vitro DNA slippage assay | | |
| SG-FS1 | CGTGTCGCCCTTC | 5' $^{32}$P-labeled primer DNA strand |
| SG-FS1 | GGG*TTTT*GAAGGGCGACACG | unlabeled template strand (4T) |
| SG-FS1 | GGG*TTTTTT*GAAGGGCGACACG | unlabeled template strand (6T) |
| SG-FS1 | GGG*TTTTTTTT*GAAGGGCGACACG | unlabeled template strand (8T) |
| SG-FS1 | CCC*AAAA*GAAGGGCGACAC | unlabeled template strand (4A) |
| SG-FS1 | CCC*AAAAAA*GAAGGGCGACAC | unlabeled template strand (6A) |
| SG-FS1 | CCC*AAAAAAAA*GAAGGGCGACAC | unlabeled template strand (8A) |

15 bp homology with linearized vectors or between two PCR fragments in bold letters.
Streptavidin tag underlined.
Catalytic dead and steric gate mutations in bold letters and underlined.
Homo-oligonucleotide runs in italic.

Table S3. Primers used in this study

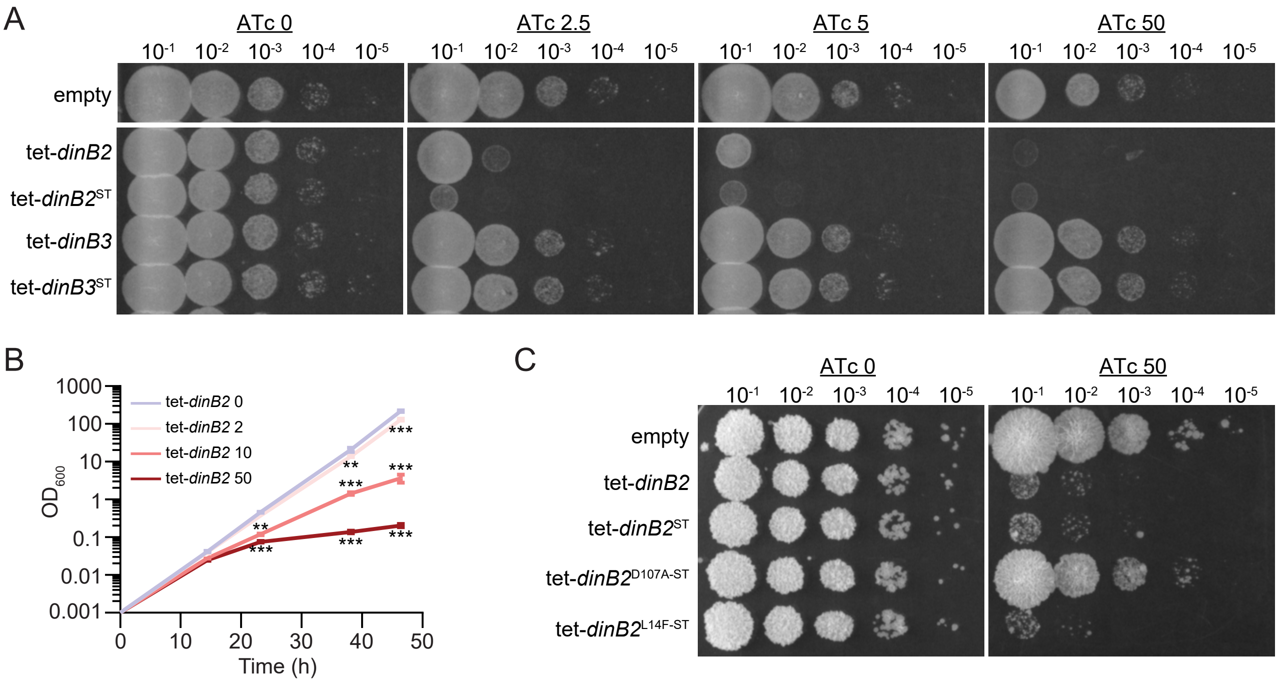


**Figure S1. DinB2 overexpression induces growth defect in *M. smegmatis*. (A)** and **(C)** Growth of indicated strains of *M. smegmatis* on agar medium containing the indicated concentrations of inducer (ATc) in agar. **(B)** Liquid growth of *M. smegmatis* carrying the *dinB2* expression plasmid in presence of the indicated concentrations of ATc. Results shown are means (± SEM) of data obtained from biological triplicates. Stars above or under the means mark a statistical difference with the reference strain (0nM of inductor) (**, P<0.01; ***, P<0.001). Empty=empty vector, tet=Atc inducible promoter, DinB2=*M. smegmatis* DinB2, DinB3=*M. smegmatis* DinB3, ST=Streptavidin tag, D107A=catalytically inactive *M. smegmatis* DinB2, L14F=Steric gate mutant of *M. smegmatis* DinB2..

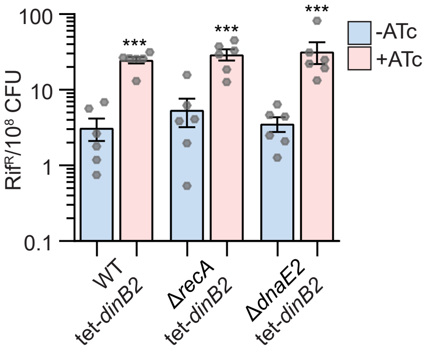


**Figure S2. Mutation frequency in Δ*recA* and Δ*dnaE2* backgrounds after DinB2 overexpression.** Rifampicin resistance (rif$^R$) frequency in indicated strains in absence (blue) or presence (red) of inducer (ATc 50 nM). Results shown are means (± SEM) of data obtained from biological replicates symbolized by grey dots. Stars above or under means mark a statistical difference with the same strain without inducer (***, $P<0.001$).

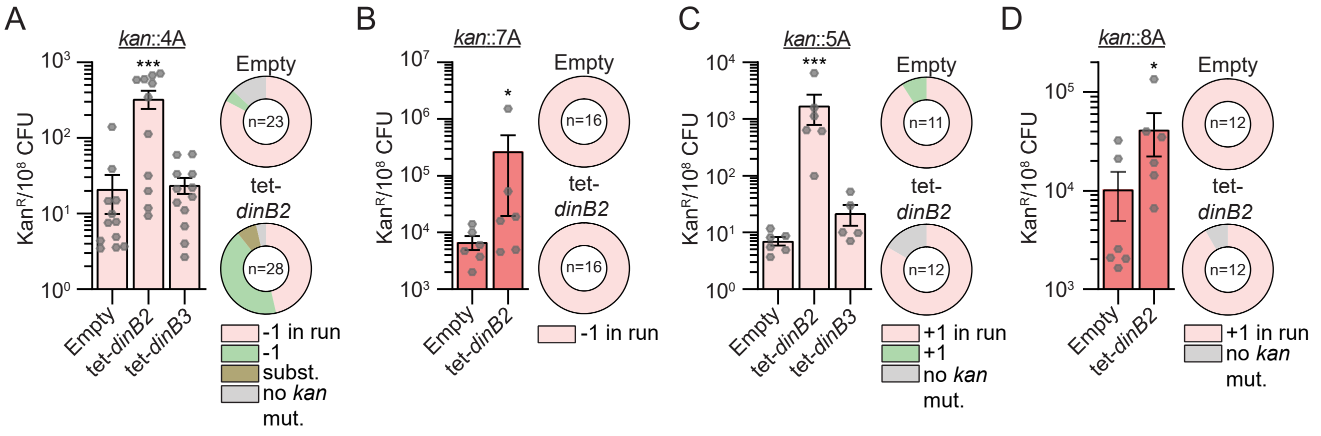


**Figure S3. -1 and +1 frameshift mutagenesis in diverse runs of A after DinB2 and DinB3 overexpression.** Kan$^R$ frequencies in the indicated strains carrying indicated mutation reporters (*kan*::4A **(A)**, *kan*::7A **(B)**, *kan*::5A **(C)**, and *kan*::8A **(D)**) in presence of ATc 50 nM. Results shown are means (± SEM) of data obtained from biological replicates symbolized by grey dots. Stars above the means mark a statistical difference with the reference strain (empty) (*, P<0.05; ***, P<0.001). Relative (pie chart) and absolute (bar chart) frequencies of nucleotide changes detected in *kan* of kan$^R$ cells represented with colors: pink=-1 or +1 frameshift in the homo-oligonucleotide run, green=-1 or +1 frameshift localized outside of the run, brown=substitution mutations, and grey=no detected mutation. The number of sequenced kan$^R$ colonies is given in the center of each pie chart. Empty=empty vector, tet=Atc inducible promoter, DinB2=*M. smegmatis* DinB2, DinB3=*M. smegmatis* DinB3.

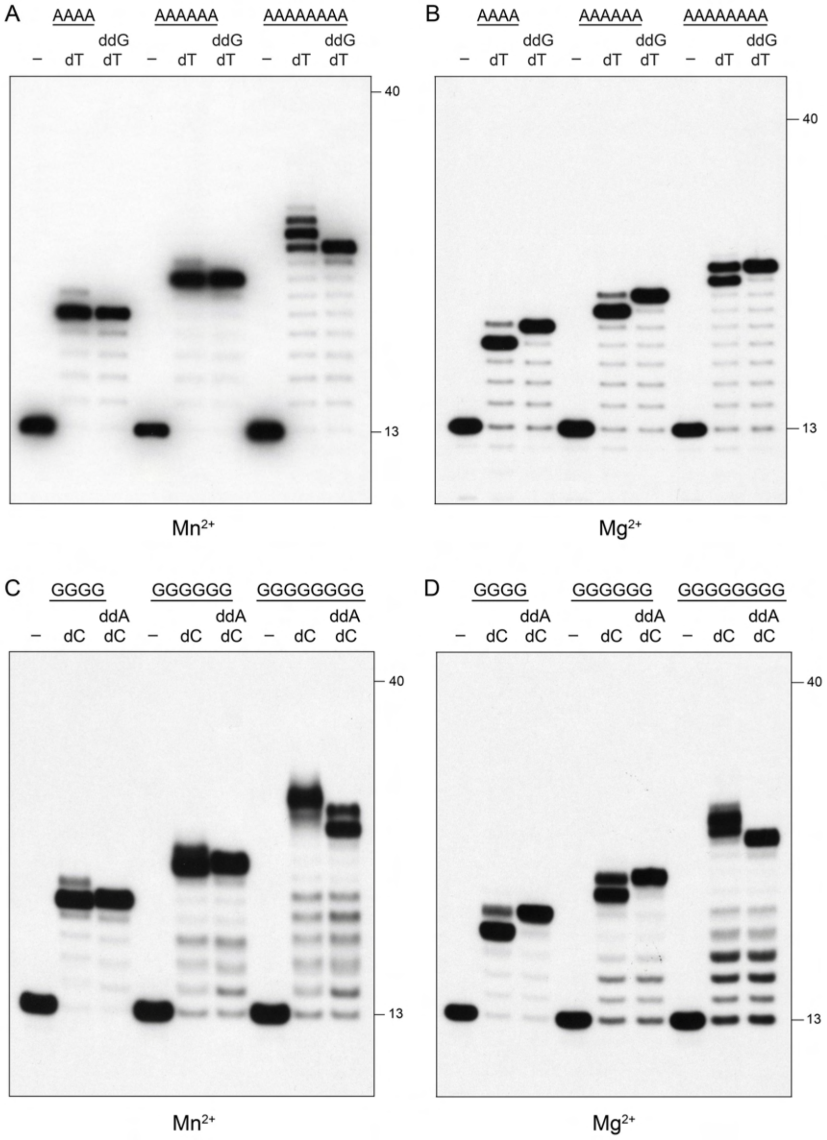


**Figure S4. Pol1 is not prone to slippage.** Reaction mixtures containing 10 mM Tris-HCl, pH 7.5, 5 mM MnCl2 or MgCl2 as specified below each panel, 1 pmol 5' $^{32}$P-labeled primer-template DNAs with A4, A6, A8, G4, G6, or G8 runs in the template strand (included as indicated above the lanes), nucleotides as specified, and 10 pmol Pol1 POL domain were incubated at 37˚C for 15 min. Pol1 was omitted from reactions in lanes –. The reaction products were analyzed by urea-PAGE and visualized by autoradiography. The positions of the 13-mer primer strand and a 5' 32P-labeled 40-mer oligonucleotide size markers analyzed in parallel are indicated on the right.

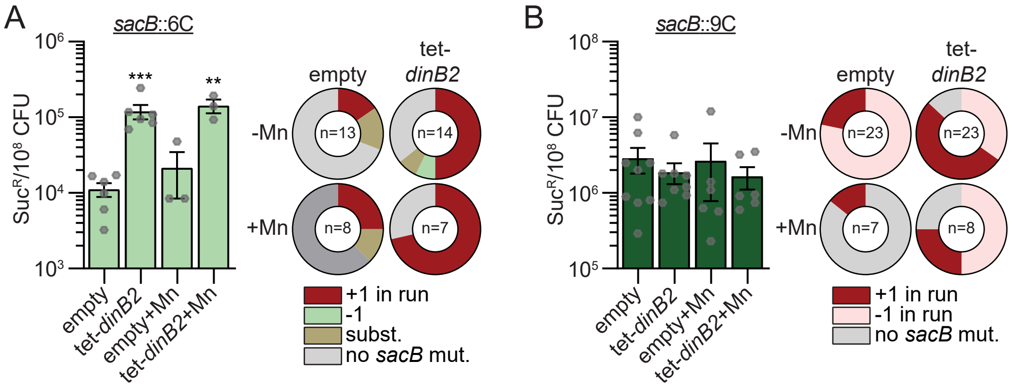


**Figure S5. DinB2 does not incorporate long slippage products in vivo.** Suc$^R$ frequencies in the indicated strains carrying indicated mutation reporters in presence of inducer and supplemented or not with 50 mg/L. Results shown are means (± SEM) of data obtained from biological replicates symbolized by grey dots. Stars above the means mark a statistical difference with the reference strain (empty) (**, P<0.01; ***, P<0.001). Relative frequencies of nucleotide changes detected in *sacB* of suc$^R$ cells are represented with colors: dark red= +1 frameshift in the homo-oligonucleotide run, light red= -1 frameshift in the homo-oligonucleotide run, green=-1 frameshift localized outside of the run, brown=substitution mutations, and grey=no detected mutation. The number of sequenced suc$^R$ colonies is given in the center of each pie chart. Empty=empty vector, tet=Atc inducible promoter, DinB2=*M. smegmatis* DinB2.
